# Hippocampal ripples predict motor learning during brief rest breaks in humans

**DOI:** 10.1101/2024.05.02.592200

**Authors:** Martin Sjøgård, Bryan Baxter, Dimitrios Mylonas, Megan Thompson, Kristi Kwok, Bailey Driscoll, Anabella Tolosa, Wen Shi, Robert Stickgold, Mark Vangel, Catherine J. Chu, Dara S. Manoach

**Author notes:** Corresponding author at: Massachusetts General Hospital, 149 13th Street CNY-2618, Charlestown, MA 02129. These authors have contributed equally to this work and share first authorship (Martin Sjøgård, Bryan Baxter). These authors have contributed equally to this work and share last authorship (Catherine J. Chu, Dara S. Manoach). Contributions: DSM, CJC, MT, and BB designed and supervised the study. BB, MT, KK, BD, and AT participated in data acquisition. MS, BB, and WS, processed electrophysiological data. MS and BB analyzed the data. MV provided statistical consultation. RS contributed to the interpretation of findings. DSM, MS, CJC, and BB wrote the manuscript. All authors contributed to the critical reading and editing of the manuscript.

## Abstract

Critical aspects of motor learning and memory happen offline, during both wake and sleep. When healthy young people learn a motor sequence task, most of their performance improvement happens not while typing, but offline, during interleaved rest breaks. In contrast, the performance of patients with dense amnesia due to hippocampal damage actually gets worse over the rest breaks and improves while typing. These findings indicate that an intact hippocampus is necessary for offline motor learning during wake, but do not specify its mechanism. Here, we studied epilepsy patients (n=17) undergoing direct intracranial EEG monitoring of the hippocampus as they learned the same motor sequence task. Like healthy young people, they showed greater speed gains across rest breaks than while typing. They also showed higher hippocampal ripple rates during these rest breaks that predicted offline gains in speed. This suggests that motor learning during brief rest breaks during wake is mediated by hippocampal ripples. These results expand our understanding of the role of hippocampal ripples beyond declarative memory to include enhancing motor procedural memory.

**Significance Statement:** In patients with epilepsy undergoing direct intracranial EEG monitoring of the hippocampus, we found a higher rate of ripples during the brief rest breaks of a motor sequence task than during task execution. These ‘offline’ hippocampal ripples predicted the amount of performance improvement over the break. We conclude that the hippocampus contributes to motor learning during brief rest breaks, and propose that this offline learning is mediated by ripples.

---

Hippocampal ripples, brief bursts of ∼70-150 Hz activity, contribute to the consolidation of hippocampally dependent memories. In rodents, specific hippocampal neuronal firing patterns during spatial navigation represent ongoing experience. During the wakeful rest and sleep that follow, these firing patterns are replayed in a temporally compressed format and coincide with hippocampal ripples.^1, 2^ Hippocampal ripple rate predicts subsequent performance and disrupting ripples during either offline state impairs performance, suggesting that ripple-related memory replay is a common mechanism of offline memory consolidation during both wake and sleep.^3-5^

Recent studies of humans also tie offline learning during wake to hippocampal function, even for a motor procedural task, which is not traditionally considered to be hippocampally dependent. When healthy young participants learn a motor procedural task (i.e., the finger tapping Motor Sequence Task, MST), most of the improvement in performance happens offline, during brief (10-30 s) rest periods between typing trials, and not during the actual typing. This phenomenon, labeled micro-offline gains, to distinguish it from the more macro-scale offline learning that occurs over hours of sleep,^6^ is associated with neuroimaging measures of increased hippocampal activation and sequential memory replay during the rest breaks that predict the level of offline improvement.^7, 8^ In contrast, patients with dense amnesia due to severe bilateral hippocampal damage, despite intact overall learning of the MST, actually show negative micro-offline gains (i.e., losses), whereas their healthy, age-matched peers retain their learning over breaks.^9^ Collectively, these findings suggest that the hippocampus is required to either improve (in young adults) or maintain (in older adults) motor memory over brief periods of wakeful rest. Here, we test the hypothesis that offline improvement of motor memory during these brief periods of wakeful rest depends specifically on hippocampal ripples. To do so, we studied patients with epilepsy undergoing invasive monitoring with hippocampal electrodes as part of a pre-surgical work-up. We divided the MST into its online (during typing) and offline (during interleaved rest breaks) components and examined ripples during each period and their relations with performance improvement (i.e., micro-online and -offline gains in speed).

## Results

Seventeen epilepsy inpatients (age: 31+/-12; range: 17-56; 5M,12F) who met all inclusion and exclusion criteria trained on the MST and were included in the analyses. The MST involves repeatedly typing a five-digit sequence (e.g., 4-1-3-2-4) “as quickly and accurately as possible” for 12 30-s trials separated by 30-s rest periods (Fig. 1A). Participants typed with the hand contralateral to the hippocampus with implanted electrodes. Typing speed was quantified as the inverse of the average interval between adjacent key presses within each correctly typed sequence (i.e., key presses/s).^6^ Micro-online gains were defined as the difference in typing speed between the first and the last correct sequence of each 30-s trial. Micro-offline gains were defined as the difference in typing speed between the last correct sequence of one trial and the first correct sequence of the next. Total gains are the sum of micro-offline and micro-online gains. The micro-online gain from the last (12th) typing trial was excluded from analyses as it is not followed by a corresponding rest period. Family-wise error rate corrected probability levels are reported to control for multiple comparisons.

**Fig. 1:**
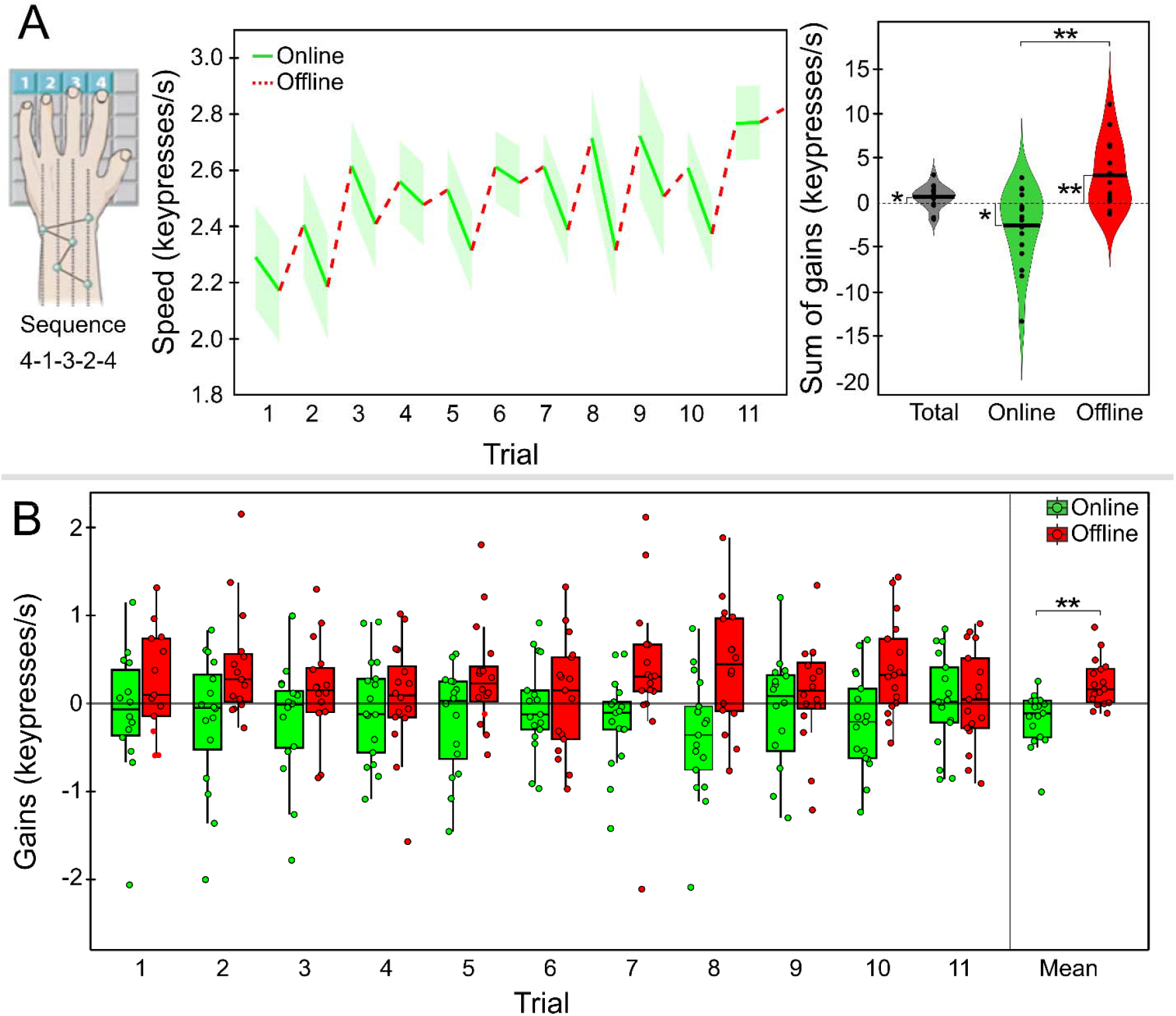
Micro-online and -offline gains across MST trials. ***A*** : *Left*: Participants rested four fingers on a key pad with keys labeled 1, 2, 3, 4. They were instructed to repeatedly type a five digit sequence (e.g., 4-1-3-2-4) “as quickly and accurately as possible.” During the 30 s typing trials, the screen was green and the sequence was displayed on top. After 30 s, the screen turned red and participants rested for 30 s. Training consisted of 12 typing trials and 11 rest breaks. *Middle*: Plot of mean online (green solid line) and offline (dashed red line) gains over the 30 s of each typing and rest period across participants (x-axis). *Right*: Violin plots showing distribution of the sum of total gains, micro-online gains and micro-offline gains across 11 trials across participants. The black dots represent participants’ means and horizontal lines represent group means. Significant differences are indicated with asterisks (**\*** = p<.05, **\*\*** = p<.005). ***B*** : Boxplots of micro-online and -offline gains by trial. Each circle is the gain for a participant in a given trial. Boxes extend to the median and 25^th^ and 75^th^ percentile of gains. Whisker lines extend to the maximum/minimum non-outlier values (less than 1.5 of the interquartile range from the upper or lower quartile). Horizontal lines represent trial medians. The rightmost boxplot pair represents the mean of micro-online and -offline gains across trials for each participant.

Participants showed significant learning over the course of MST training (Total gains: 0.5±0.06; t_16_=3.41, p=.004) and most of this improvement happened offline (Fig. 1). Participants showed significant micro-offline gains (2.3±2.6; t_16_=3.69, p=.002) and lost speed online, while typing (i.e., negative micro-online gains: -1.9±3.0; t_16_=-2.59, p=.02). Across participants, summed micro-offline gains were significantly greater than micro-online gains (4.2±5.6; t_16_=3.13, p=.006; Fig. 1A, right), and median micro-offline gains were numerically greater than micro-online gains on every trial (Fig. 1B). The finding of greater learning during offline (rest) than online (typing) periods is consistent with previous studies of healthy young adults.^6, 7^

We identified hippocampal ripples (70-150 Hz) during MST training using an automated ripple detector adapted from previous studies.^10-12^ Ripples were detected in intracranial EEG recordings targeting the hippocampus contralateral to the hand that performed the MST. Detected ripples showed the characteristic sharp-wave and oscillatory ripple band activity in the wideband filtered signal (0.3-250 Hz) during both online and offline periods (Fig. 2A; see Supplementary Fig. S1 for additional examples of detected ripples).^13^ The mean frequency, amplitude, duration, and number of cycles of detected ripples did not significantly differ between online and offline periods (Fig. 2B). We calculated the ripple rate (ripples/s) for each participant during each online and offline period. Across participants and trials, the mean ripple rate was higher during offline than online periods (Fig. 3A; 0.17±0.10 vs. 0.11±0.04; t_16_=2.27, p=.038). Median ripple rate was also numerically higher during offline than online periods on every trial.

**Fig. 2:**
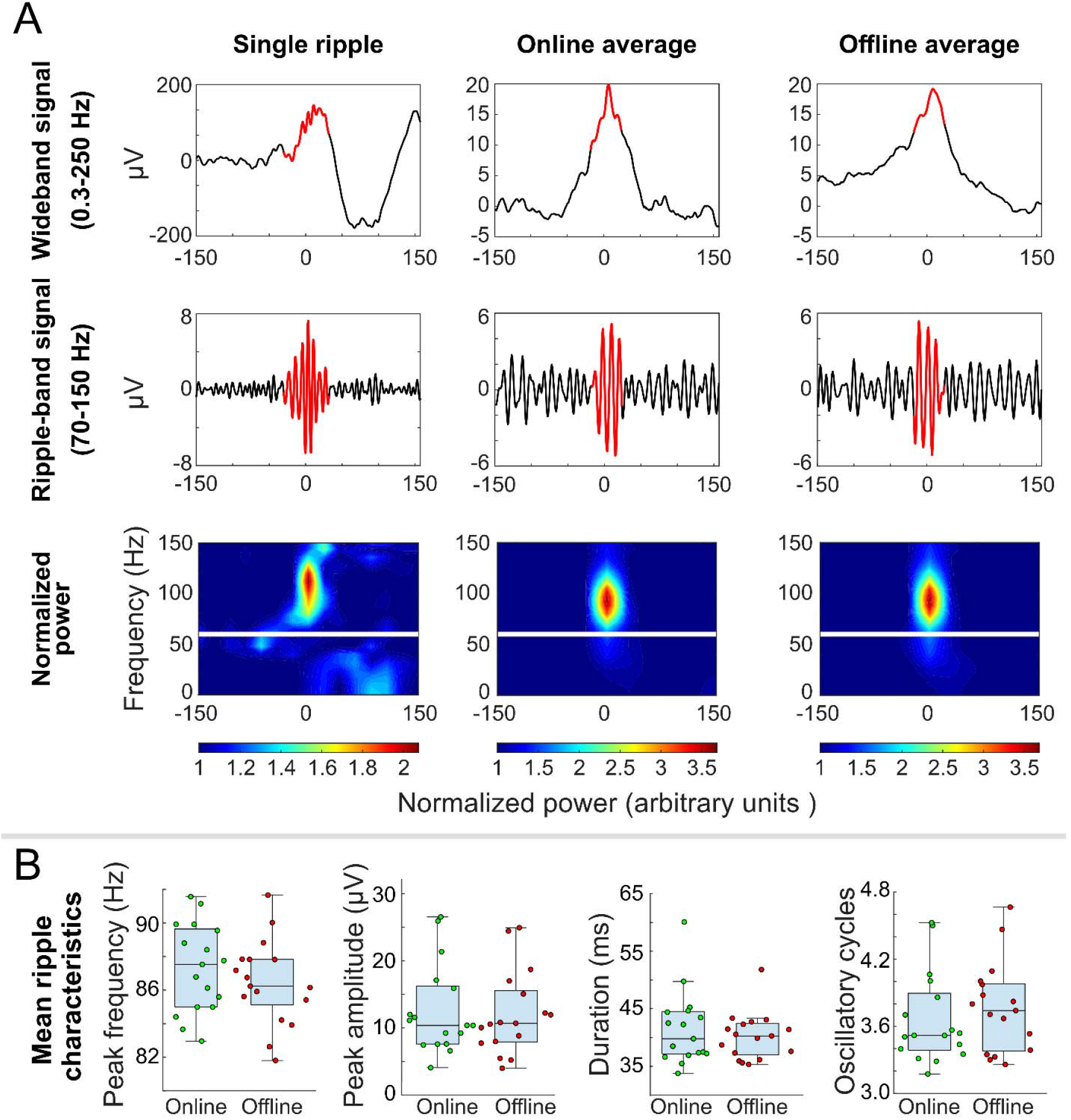
Oscillatory and spectral characteristics of detected ripples. ***A***: Wide-band filtered signal, 70-150 Hz filtered signal, and time frequency spectrograms for a single ripple and averaged ripples (highlighted in red) during online (typing) and offline (rest) periods. Ripple peaks were aligned before averaging. Time frequency spectrograms are normalized to the mean across the entire MST run at each frequency. The white line indicates where 60 Hz signal was removed using a notch filter. Each plot is centered on maximum power in the 70-150 Hz filtered signal at the time of a detected ripple. ***B***: Online and offline ripple characteristics. Each dot represents the mean value of a participant’s ripples’ peak frequency, peak amplitude, duration, and number of oscillatory cycles. Boxes extend to the median and 25^th^ and 75^th^ percentile. Whisker lines extend to the maximum/minimum non-outlier values (less than 1.5 of the interquartile range from the upper or lower quartile). Horizontal lines represent medians.

**Fig. 3:**
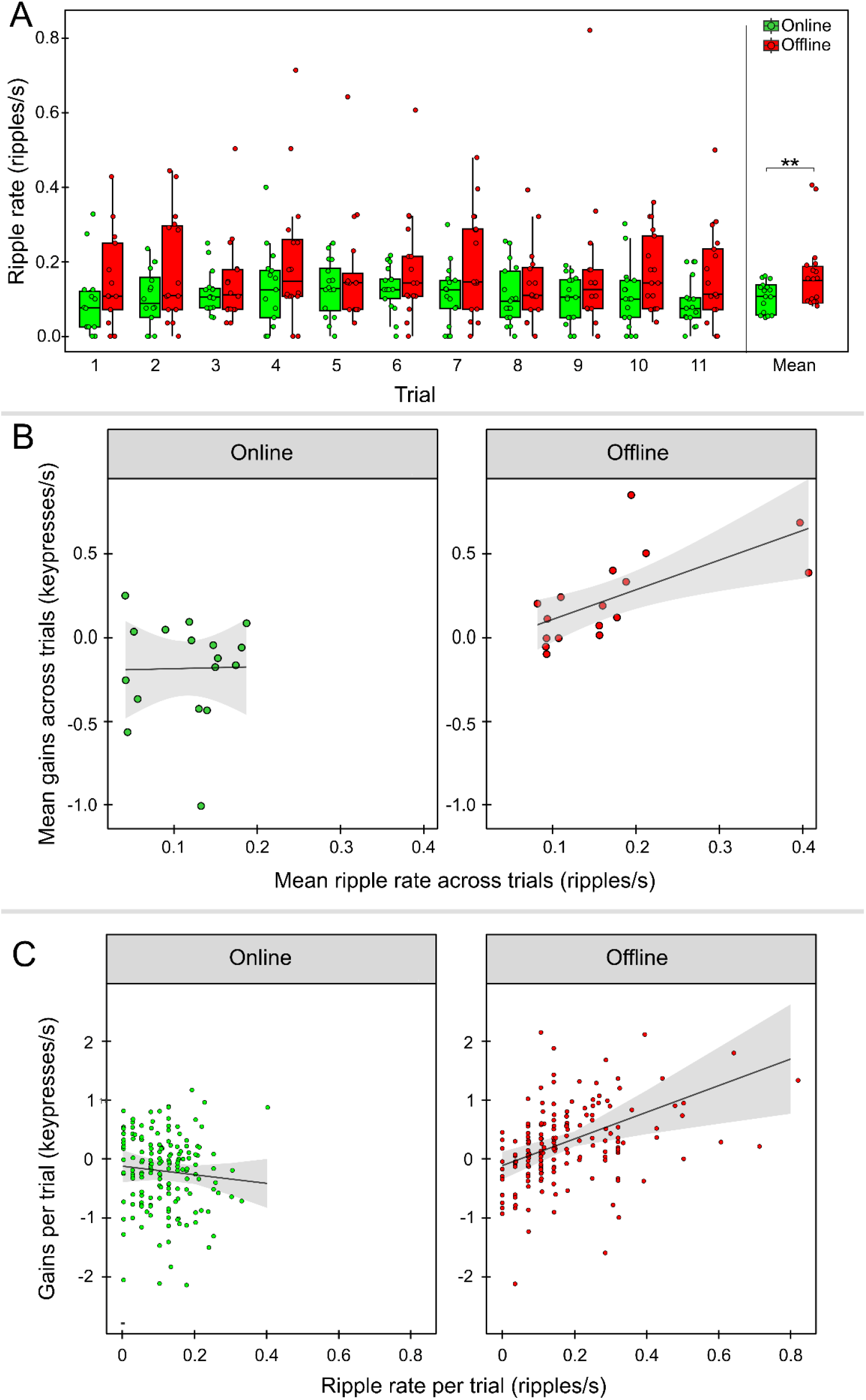
Hippocampal ripple rates during online and offline periods and their relations to micro-online and -offline gains. ***A*** : Boxplots of hippocampal ripples during online and offline periods for each trial. Each circle is an individual data point. Boxes extend to the median and 25^th^ and 75^th^ percentile of ripple rate across participants. Whisker lines extend to the maximum/minimum non-outlier values (less than 1.5 of the interquartile range from the upper or lower quartile). Horizontal lines represent medians. Mean online and offline ripple rates across trials per participant (far right). ***B*** : Scatter plots of mean ripple rate and mean gains across trials. Each participant is a circle. The shading represents the standard error of the regression line. ***C*** : Scatter plots of ripple rates and gains per trial for each participant for online and offline periods.

Participants with a higher ripple rate during the rest breaks showed correspondingly greater offline gains (Fig. 3B right; r=.64, p=.005), consistent with our hypothesis that offline memory consolidation during wake depends on hippocampal ripples. In contrast, the ripple rate during typing did not correlate with total online gains (Fig. 3B left; r=.006, p=.98). We then investigated whether ripple rate predicted gains on a trial-by-trial basis (see data in 1B and 3A). Ripple rate significantly predicted individual trial gains (β est.=2.38; 95% CI: 1.13—3.64; p=.0008) and this relation differed for online vs. offline periods (β est.=-3.11; 95% CI: -4.68— - 1.54; p=.0001). Post-hoc tests found a significant effect of offline ripple rate on offline gains (β est.=2.26; 95% CI: 0.84—3.68; p=.002) but no effect of online ripple rate on online gains (β est.=-0.73; 95% CI: -2.56—1.10; p=.4; Fig. 3C).

## Discussion

Our primary findings are that patients with epilepsy undergoing invasive EEG monitoring of the hippocampus *i)* have more hippocampal ripples during brief, 30-s breaks while learning a motor procedural task than while actually performing the task and *ii)* that these offline ripple rates predict task improvement across these same offline periods, both on a subject-by-subject and a trial-by-trial basis. We did not observe these relationships between the online ripple rate and performance changes across typing periods. These findings are consistent with rodent studies showing ripple-related memory replay during the wakeful rest that follows spatial navigation.^14, 15^ Disrupting these ripples impairs memory, suggesting that they play a causal role.^3, 4^ Our findings also complement human neuroimaging findings of hippocampal activation and sequential memory replay during MST rest breaks that predict the level of offline improvement.^7, 8^ The absence of offline learning during both wake^9^ and sleep^16^ in patients with dense amnesia due to hippocampal damage, together with the present findings, suggests that the hippocampus is necessary for offline learning and that at least during wake, this learning depends on ripples. These findings support the possibility that hippocampal ripples are a common mechanism of motor memory consolidation during wake and sleep in humans.

Procedural memory is classically thought not to involve the hippocampus, but instead to rely on the striatum, based on findings that motor procedural performance can improve despite bilateral hippocampal damage.^17-19^ How the striatum and hippocampus might interact to mediate motor learning is unclear, with some evidence suggesting competitive interactions (e.g.,^20, 21^). More recently, distinctions between offline and online motor learning have emerged. Human neuroimaging studies show hippocampal activation while learning procedural motor tasks,^22, 23^ including the MST, and more so during rest than typing.^7, 8^ But neuroimaging studies cannot address questions of whether the hippocampus is simply engaged during offline motor learning or whether it contributes to that learning, and if the latter, by what mechanism? Our studies of people with amnesia due to hippocampal damage demonstrate that the hippocampus is necessary for the offline consolidation of motor procedural learning over both sleep^16^ and wake.^9^ The present study supports a critical role for hippocampal ripples in the wakeful offline consolidation of motor procedural learning. The findings complement and provide a potential mechanistic explanation for a growing body of neuroimaging findings of hippocampal engagement during wakeful rest following motor learning (e.g., ^7, 8, 24, 25^). This body of work expands our understanding of the role of the hippocampus beyond declarative memory to include the formation of motor procedural memory.

There were significantly fewer ripples during typing than rest periods, and no correlation between online ripple rates and online gains. This likely reflects differences in hippocampal activity during active vs. offline learning, as is seen in rodent studies.^26^ During spatial navigation, hippocampal place cells exhibit location-specific firing, primarily in the theta band, to provide a map of the environment (a similar phenomenon is seen in humans^27^). During the wakeful rest that follows, hippocampal neurons repeatedly reactivate recent experiences via the firing of place cells on a much faster time scale during ripples.^14, 15^ This ripple-related replay correlates with memory improvement^28^ and disrupting these ripples consistently impairs memory, consistent with a causal role.^3, 4^ Indirect evidence from human neuroimaging studies also links hippocampal ripples to memory reactivation^29^ and hippocampal sequential memory replay to memory improvement during rest breaks of the MST.^8^ Although the present study does not measure memory replay, given the rodent and human studies linking ripples to replay, we speculate that the increased ripple rate we observed during rest breaks and its correlation with performance improvement reflect the occurrence of ripple-related memory replay. Further, the loss of ripples and ripple-related memory replay may be the culprit in the losses of offline motor sequence learning during rest breaks and sleep in amnesia.^9, 16^

In the present study, as in previous studies of young adults, participants showed online losses of speed followed by marked offline improvements.^6^ This pattern could reflect motor fatigue accumulating during typing and its dissipation during the rest break that follows.^30^ If this were the case, any learning acquired during the previous typing period could only be fully expressed after rest. Our data, however, indicate that the motor slowing seen during typing periods is unlikely to be due solely to muscle fatigue. When we compare typing speed on the screening task, which did not involve learning (2 30-s trials of typing 1-2-3-4 separated by a 30-s rest period) to the first two MST trials in the 7 subjects for whom we had this data, participants typed almost twice as fast on screening (4.1±2.9 vs. 2.2±1.3 keypresses/s; t(6)=2.8, p=.03), but showed less than half the online losses (−0.08±0.4 vs. -0.22±0.4 keypresses/s; t(6)=2.3, p=.05). These data are incompatible with micro-offline gains on the MST representing recovery from muscle fatigue since the same subjects proved themselves capable of sustaining higher typing speeds on an over-learned sequence across 30-s trials. More likely, they reflect recovery from psychomotor fatigue. Alternatively, this pattern could reflect an enhancement of learning during rest as a result of ripples. These explanations are not mutually exclusive, though the correlations of micro-offline gains with hippocampal ripples, both across participants and across trials, argues for ripples as an important driver of offline learning. So while we cannot definitively conclude that ripples mediate motor memory improvement (i.e., learning), we can reasonably conclude that our findings provide evidence that ripples mediate the offline consolidation of motor memory during wake, whether referred to as improvement, recovery, stabilization, or maintenance.

In conclusion, the hippocampus is necessary for the offline consolidation of motor learning during both brief rest breaks and sleep and this offline learning during wake appears to be mediated by ripples.

## METHODS

### Participants

Inpatients with epilepsy undergoing continuous intracranial EEG monitoring of the hippocampus as part of their clinical care were evaluated for participation. Twenty-three participants who met the following inclusion criteria enrolled in the study: ≥12 years old; no prior cortical or subcortical resection; no prior neurosurgical procedure that was expected to interfere with sleep oscillations; estimated IQ ≥70 based on neuropsychological assessment and/or the single word reading subtest of the Wide Range Achievement Test-4 (WRAT-4^31^); no history of marked developmental delay or marked motor impairment. Participation for three participants was discontinued after they failed to pass a motor screening test requiring them to correctly type the sequence “1,2,3,4” a total of 24 times during two 30 s trials separated by a 30 s rest period. Of the 20 remaining participants, two were excluded because invalid trials on the Motor Sequence Task (MST) affected the calculation of gains in ≥7 of the 22 online and offline periods (see Excluded MST Data below), and one participant was excluded due to excessive amounts of gamma noise in hippocampal recordings (see Signal Preprocessing below). The final sample was comprised of 17 participants (age: 31+/-12; range: 17-56; 5M,12F; see Table S1 for additional participant information). All participants provided informed consent in accordance with the Partners Institutional Review Board and the Declaration of Helsinki.

### Motor Sequence Task (MST)

#### MST Hand Selection

Participants performed the MST with the hand contralateral to the hippocampus with implanted electrodes (n=9). If participants had bilateral hippocampal implants, they used the hand contralateral to the presumed healthier hippocampus (n=8) based on review of medical records, MRI, and hippocampal electrophysiology by a board-certified epileptologist and neurophysiologist (CJC).

#### MST Warmup

To acclimate participants to the structure of the MST, they were administered a ‘warmup’ task prior to MST training. Participants rested the index and middle fingers of the hand that they were not using for the MST on the keypad buttons labeled 3 and 4. They were instructed to repeatedly type the sequence 3-4 “as quickly and accurately as possible” for three 10 s typing periods separated by 10 s rest breaks. The sequence remained on the screen for the duration of the task. During the typing periods the screen remained green and a dot appeared in a horizontal line under the sequence with each key press. After the line reached the right border, the dots disappeared one at a time, from right to left, with each additional key press. After 10 s, the screen turned red, and participants rested for 10 s. A countdown of the number of seconds until the screen turned green was displayed as spelled out numbers. The last three numbers were replaced with tones to alert the participants to get ready to resume typing when the screen turned green again.

#### MST Administration

The MST had the same structure as the warmup with the following exceptions: the participant used four fingers on the keypad (index, middle, ring, pinkie) to type a 5-digit sequence (e.g., 4-1-3-2-4) and there were twelve 30 s typing trials separated by 30 s rest periods. If a participant did not follow task instructions (e.g., continuously used the incorrect finger for a key), they were corrected in real time by the experimenter.

#### MST outcome measures

The primary outcome measures were total gains, micro-online gains, and micro-offline gains in typing speed. Typing speed for each correctly typed sequence was quantified as the inverse of the interval between the first and last keypress of the sequence (i.e., key presses/s). Micro-online gains are defined as the difference in typing speed between the first and the last correct sequence of each typing trial.^6^ Micro-offline gains are defined as the difference in typing speed between the last correct sequence of a trial and the first correct sequence of the next trial. Total gains are the sum of micro-offline and micro-online gains per trial and are equal to the difference in typing speed between the first correct sequence of one trial and the first correct sequence of the next. The micro-online gain from the last (12^th^) typing trial was not included in the calculations as there is no subsequent rest period, leaving a total of 11 online and 11 offline gains per participant.

#### Excluded MST data

It is not possible to calculate online gains on trials with no correctly typed sequences, nor is it possible to calculate offline gains for the rest period before and after it. The same is true for trials that were rendered invalid for other reasons, including interruptions by clinical staff or a participant’s failure to follow task instructions (e.g., using the same finger on more than one key). The data from the two participants were excluded from analysis because invalid trials affected the calculation of gains for ≥7 of the 22 online and offline periods. In included participants, the gains of an average of <1 period per participant could not be scored (4% of the periods).

### Hippocampal Ripple Measurement

#### Electrode selection

All participants had intracranial recordings targeting the hippocampus. In two participants, foramen ovale electrodes were used. In four participants, a single stereotactically placed electroencephalography (SEEG) electrode was placed that targeted the hippocampal body, and this electrode was used for analysis. In the remaining 13 participants, an SEEG electrode targeting the hippocampal head was used for analysis. In each SEEG case, the three deepest contacts were bipolar referenced, creating two channels per subject for ripple detection.

#### Data acquisition

Data were acquired at 1024 or 2048 Hz with a Natus Quantum or EMU40 clinical system (Natus Medical Inc., Middleton, WI, USA). Data recorded at 2048 Hz were downsampled to 1024 Hz.

#### Signal preprocessing

Data were high pass filtered above 0.1 Hz using a two-pass Finite Impulse Response (FIR) filter in Matlab using the Fieldtrip toolbox.^32^ Sixty Hz line noise and its first harmonic were removed using a two-pass bandstop Butterworth filter. All signals were visually inspected and high-amplitude artifacts were removed from four participants based on visually determined subject-specific thresholds. Epileptiform spikes were detected using the Reveal algorithm in Persyst 14 software with automatic thresholding (Persyst Development Corporation, Sand Diego, CA, USA).^33^ High-amplitude spikes missed by the algorithm were manually marked based on visually determined subject-specific thresholds, and all spikes (peak ± 35 ms), were ignored in analyses.^34^ To avoid using channels with excessive high frequency noise within the ripple-band, we identified those that did not have the expected steep fall-off of power as a function of frequency.^35^ In each selected channel, we computed the linear fit between the log power and the log frequency. If the slope of this fit was not lower than -2, we used the closest channel that met this criterion (2 participants). One participant’s data were excluded from analyses since none of the channels met this criterion.

#### Automated Ripple Detection

For each participant, ripples were detected across all online and offline periods using a common amplitude threshold. Hippocampal electrodes were bipolar referenced to adjacent electrodes on the same lead. The bipolar signal was bandpass filtered (70-150 Hz) and the envelope of the signal was calculated using the Hilbert transform. The root-mean-squared amplitude of the ripple band envelope was calculated for each electrode. The threshold for ripple detection was set to the upper 95^th^ percentile, which is in the range of other studies.^11, 12^ (Note that the findings are qualitatively the same using the upper 99^th^ percentile as a threshold (see Fig. S2)).The threshold was calculated using the full signal with artifacts and spikes removed. Each detected ripple was required to exceed this threshold for >6 ms and to have at least 6 peaks, indicating 3 full cycles, in both the raw and the bandpass filtered signal^36^ and have a ripple-band envelope >3 SDs above the mean during the 200 ms window around the ripple peak. If a detected ripple was within 50 ms before or after the peak of a detected spike, or if it overlapped with detected artifacts, it was rejected. To confirm that the automated detector worked as expected, an expert (CJC) visually validated 20% of both the detected online and offline ripples. The false positive rate was 4.3% and 4.1% for online and offline ripples, respectively.

#### Statistical Analyses

To determine whether online, offline, and total gains differed from zero, we used one sample t-tests. Paired t-tests were used to compare offline vs. online gains and offline vs online ripples. We also used paired t-tests to conduct control analyses of online and offline ripple characteristics (amplitude, frequency, duration, and number of cycles). To determine whether participants with higher ripple rates also had greater offline gains, we used Pearson correlations. These analyses were corrected for multiple comparisons (family-wise error rate) using non-parametric permutation testing.^37^ For each permutation, we randomly swapped half the pair labels (t-tests) or randomly shuffled the values of one variable (correlations). The null distribution for each test category was based on 10,000 permutations. In each permutation we extracted the maximum absolute test statistic within each category. A two-tailed significance threshold of p<.05 was set at the 95th percentile of each resulting distribution (absolute value thresholds: paired t-tests: t=1.77; one-sample t-tests: t=2.12; correlation coefficients: r=.47). Corrected p-values were estimated from these distributions. To assess the relation of gains with ripples on a trial-by-trial basis within participants, we employed linear mixed effects models with Subject as a random effect to account for repeated measures. We used Ripple Rate, Period (Online, Offline) and their interaction as fixed effects to predict Gains, while allowing the Intercept and Ripple Rate to vary randomly within Subject (Subject Gains ∼ Ripple Rate * Period + (1 + Ripple Rate | Subject)). To test our main hypothesis that offline ripples predict gains during offline periods, we investigated online and offline periods in separate linear models (e.g., Offline Gains ∼ Offline Ripple Rate + (1 + Offline Ripple Rate | Subject)). P-values for the mixed effect model estimates were calculated using the Kenward-Roger method.^38^ Confidence intervals of the estimates were calculated using a semiparametric bootstrap approach (5,000 iterations) implemented in R using the lmeresampler package.^39^ All models and statistical inference methods were implemented in R using the lme4,^40^ sjPlot^41^ packages.

