## Supplemental figures for "Hippocampal ripples predict motor learning during brief rest breaks in humans"

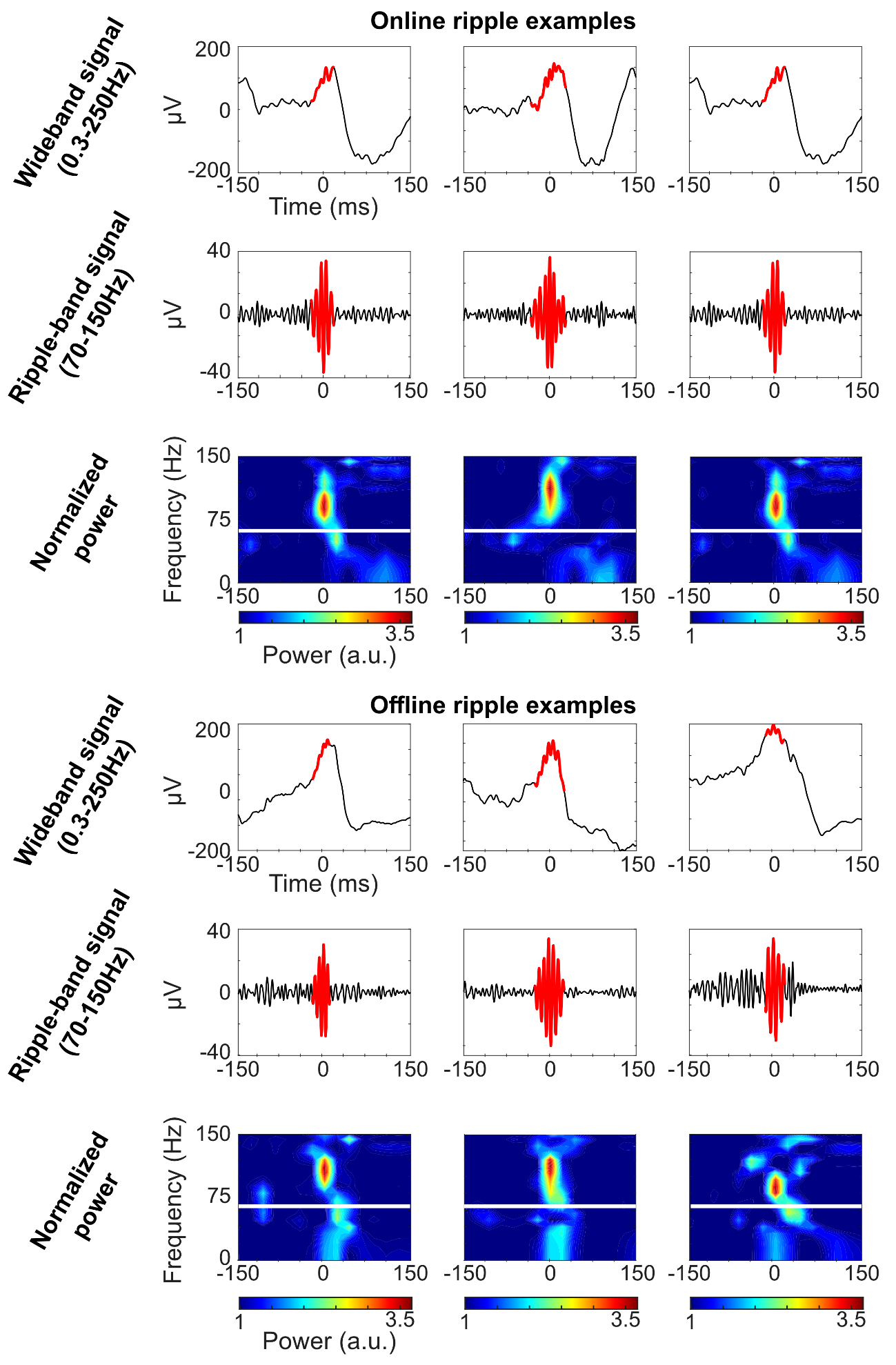

**Fig. S1: Examples of detected hippocampal ripples during online and offline periods**

Wide-band filtered signal, 70-150 Hz filtered signal, and time frequency spectrograms for three example ripples (highlighted in red) from the online (top three rows) and offline (bottom three rows) periods. Ripple peaks were aligned before averaging. Time frequency spectrograms are normalized to the mean across the entire MST run at each frequency. The white line indicates where 60 Hz signal was removed using a notch filter. Each plot is centered on maximum power in the 70-150 Hz filtered signal at the time of a detected ripple.

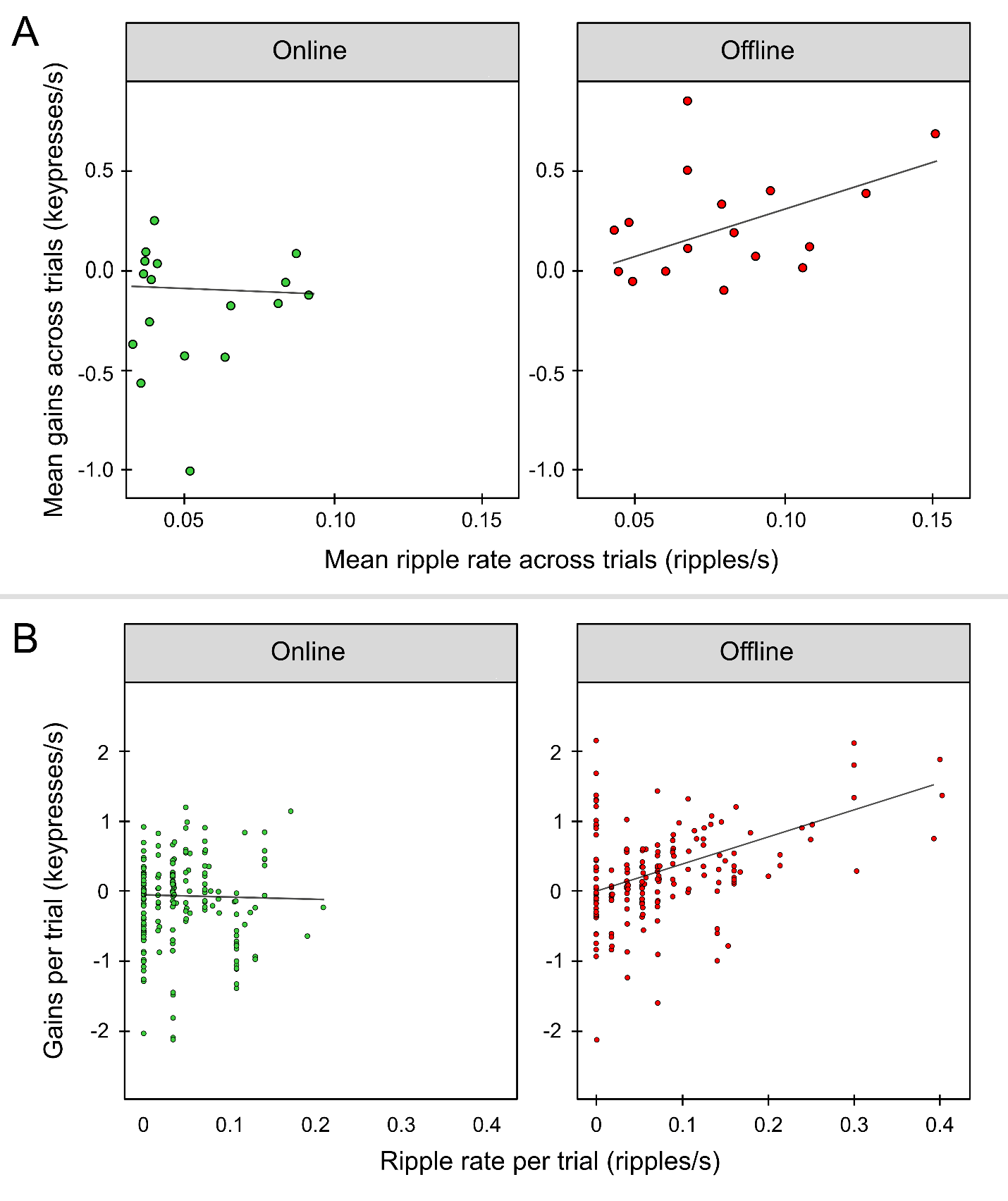

**Fig. S2: Relations of hippocampal ripples detected with the upper 99^th^ percentile threshold with micro-online and -offline gains.**

***A***: Scatter plots of mean ripple rate and mean gains across trials. Each participant is a circle. The shading represents the standard error of the regression line. Participants with a higher offline ripple rate showed correspondingly greater offline gains (right; r=.57, p=.009), whereas online ripple rate did not correlate with online gains (r=-.09, p=.78).

***B***: Scatter plots of mean ripple rate and mean gains per trial for each participant during online and offline periods. Overall ripple rate predicted individual trial gains (est.=2.02; 95% CI: 1.01—3.4; p=.002) and this relation differed for online vs. offline periods (est.= -2.98; 95% CI: -4.22— -1.98; p=.001). Post-hoc tests found a significant effect of offline ripple rate on offline gains (est.=2.02; 95% CI: 0.78—3.95; p=.007) but no effect of online ripple rate on online gains (est.=--0.33; 95% CI: -1.98—1.65; p=.55).
